## Supplementary for "Nrf1 is not a direct target gene of SREBP1, albeit both are integrated into the rapamycin-responsive regulatory network in human hepatoma cells"


#### Figure S1

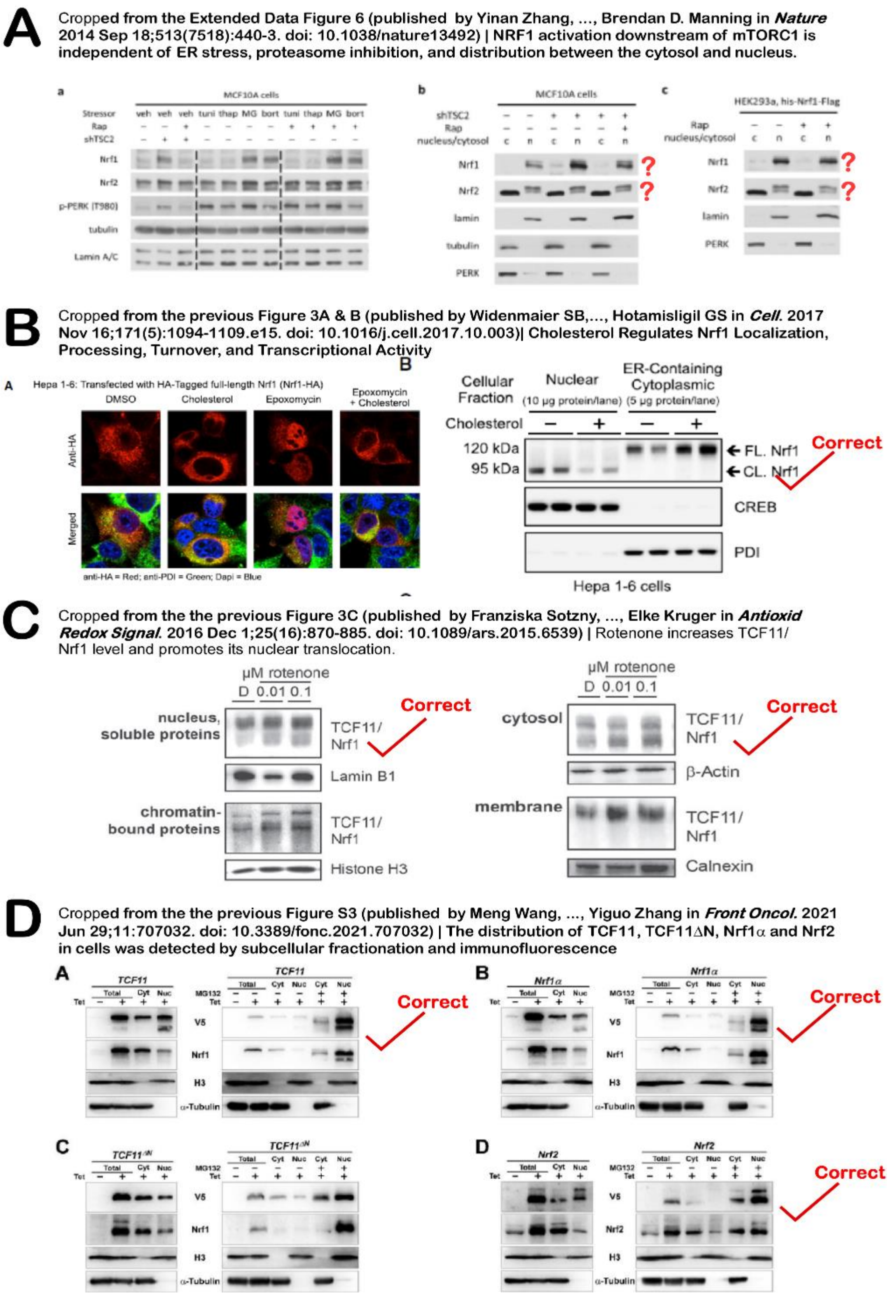


**Figure S1.** All the images are cropped from the original publications as indicated here. The relevant contents were also described in the main texts of this paper.

### Figure S2

##### *SRE* sites existing in the human *Nrf1* gene promoter regions

```
                                             TSS1
GGAGAAACTGTTCCTAGGGTGTTTGCGGCGGGCGGGTTTCCGAGTGCGGTAGCAGGAGGGGGCGGG
GAGGTAAGCGGAGGCTCCGAGCTCTAGGCCGGCCGGCGGTGGCGGCGGCGAGGCCGGGACTCGGGC
TTAGGGCCTGCTGTGGAGGCAGCGGCGGACGCCGAGCTAAGCAGTGGGTATCGGGCACATTCCTTT
CCCAGAAGGGAGGGTTGCAGCGCCGCGGGGCCGAGGCTTATGGGGCGACGGGATGGTGCTGAGGTG
CAGGAGGCGGGGGCCGAGCGGGGTTCCCGCCAGAGCCCGGCCCTGGGCCTGACAGGGAGGGGCCCC
GCGCGGGGCCGAGAGGAAAGCTGGCTCCGGGGTGGGATGGACTGGGCAGGGTTGAGGGCACGAGAA
GGCGTGGTGTGAGGCCTGGACTTGGGAGTGGGGTGGGGCCATCGGGACGGAGGGCGGGGTGGGGTT
                                   TSS2
TGCTGTAAGGGTGGGTCAAGGCCAGGGAAGTAGCACTTGTTCGCTGGCCGCCCCTGGAGGCTAGAA
GCTCCGGCGCCGAGAGTGGGCATGGCGACTTGGTCTCAGCCGGACTCGGGTATCCGGGCAGGGTGG
GACCAGCCTCAGGCTATGGGCCCTGCACCTACTTGATTTTTAAGGGACTTTGTCCGGTTCCTCGAA
CTTAATTGGCTCCTGGGAGGTTTTAGTAATAAATTGCGTGAGAACGGGAGGGTAGCGTAAGCAGGA
ATTCTGGGTCGCAATTGTGTTTACTCTGACCAGAGAGAGGGAGTGTGAGTGTGTGTGTGTGCGCAC
GTTCGTGTGTGTACGTATAGCTCGCTCTCCCCCTGTAGAATGAAGTTGGTAGATGGGTCACAACGC
AGGAGAGGAGAGGCGGTATGTCTTGGTTTGTGACATTAATGGAAGCTCAAAAGTGGAGACCAGAAA
CAGATTTACTTCCTCTGCACTGGGCAGTCTGGCTAGAGAACGTGGCCATATCCTTCTGCTTTGGCT
CAGAAGGTCATATGAATGAAAATAAAACCTGCCTATGCAAATTCTGGACGGAATAGTCCTGCGTAG
TAGGGCCCCCACCATCCCTCTCCTTTTTTTCTTTCCTTTTCTTTTCTTTCTTTTTTTTCTTTTTTT
GAGACGGAGTTTTGCTCTTGTTGCCCAGGCTGGAGTGCAGTGGCGCGATATCGGCCCACCACAACC
TCCGTTTCCTGGGTTCAAGCGATTCTCCTGTCTCAGCCTCCCGAGTAGCTGGGATTACAGGCATGC
GCCACCACGCCCGGCTAATTTTTGTATTTTTAGTTGAGATGGGGTTTCTCCATGTTGTTCAGGCTG
GTCTCAAACTCCCGACCTCGGGTGATCCGCCCTCCTCGGCCTCCCGAAGTGCTGGGATTACAGGCG
TCGGCCCTTTTCCTTTTTTTTCTTTTGTAAGTTGTGAGGATAGGAAAGGGAAATCAGATTTCAAAG
TTCAGAAAACAAAAGGCATTCTTCTTCTGGGTGAAGTGGTGGCCTGAACATTGTCCTTTTGTCCTT
CCACACTCTCACTGTTTAATAATCTTGCCTTTTTGAACTGAGAAAAGACAACTGTTATGGGTGGTT
TTCTCCCGAGGGGTGAAAATCCAGCTTTACTTTTAAGACAGGAAATGGGCTTCTCATGTGAAAATA
GAAGCTGGGTAATGCGCCTGTAGTGGGGGAGGAGGGCTGGTAAATAAAACAAAGGTGTAAGATATT
TTGGATGACCTGGTTATTGTCTGCTTATTCAAACCAAGCTACATTTTAGGATATTGCACATTTAAT
ATTCAGTAACTGAGATCATTCGTGATCAGTAGCTTTCTTTGATAGTTCTTCTATGATCGGTTGCTA
TATTTTGGGTTTTTTTTCTGAGAGTGTAAGTAGTCCTTTCTTCCTGTAAAAGTGGCCTTTAGGCCA
GGCACAGTGGCTCACGCCTGTGATCCCAGCACTTTGGGGGGCCGAGGCGGGCAGATCACCTGAGGT
CAGGAGCTTGAGACCAGCCTGGCCAACATAGTGAAACCCCATCTCTACTAAAAATACAAAAATTAG
CAGGGCGTGGTGGCAGGCGCCTGTAATCCCAGATACTTGGGAGGCTGAGGCAGGAGAATCGCTTGA
ATGCAGAAGGTGGAGGTTGCAGTGAGCACAGATCACGCCACTGCACTCTAGCCTGGGCGACAGAGT
GAGACTCTGTCTCAAAAAAAAAAAAAAGTGAAAGTGTCCTTTGAATGTGTTTTCTACCTTCTACTGT
GAAGGTTTTCCTACCCTGATCATGATGTGAAAAGTATGGAATCTTATCCTGTTGCTTTTGTGATTC
CCATTTCAGGTTTCTCTGGAAACCCCCCTGGTAAGTGTGGAGGAGGCGGGACACTCTGACCCAAGA
CGAAAGGCCTGTAGCTCCAGCCAAAGAAAATAAACCTTAGGAGGGAGAAGGAAAAAAAAAATCCAT
CAGCTGTTCCTGAGAACAGCCTGCATTGGAATCTACAGAGAGGACAACTAATGTGAGTGAGGAAGT
GACTGTATGTGGACTGTGGAGAAAGTAAGTCACGTGGGCCCTTGAGGACCTGGACTGGGTTAGGAA
CAGTTGTACTTTCAGAGGTGAGGTGTCGAGAAGGGAAAGTGAATGTGGTCTGGAGTGTGTCCTTGG
CCTTGGCTCCACAGGGTGTGCTTTCCTCTGGGGCCGTCAGGGAGCTCATCCCTTGTGTTCTGCCAG
GGTGGGGTACGGGGTTTGACACTGAGGAGGGTAACCTGCTGGCTGGAGCGGCAGAGCAGTGGCCTT
                                                               TIS
GATTTGTCTTTTGGAAGATTTTAAAAACCAAAAAGCATAAACATTCTGGTCCTTCAGCAATGCTTT
CTCTGAAGAAATACTTAACGGAAGGACTTCTCCAGTTCACCATTCTGCTGAGTTTGATTGGGGTAC
GGGTGGACGTGGATACTTACCTGACCTCACAGCTTCCCCCACTCCGGGAGATCATCCTGGGGCCCA
GTTCTGCCTATACTCAGACCCAGTTCCACAACCTGAGGAATACCTTGGATGGCTATGGTATCCACC
```

**Figure S2.** Several sterol regulatory element (SRE) consensus sites existing in the promoter of human *Nrf1* gene. Two distinct transcription start sites (TSS1 and TSS2), along with translation initiation signal (TIS) are indicated by arrows in the nucleotide locations

#### Figure S3

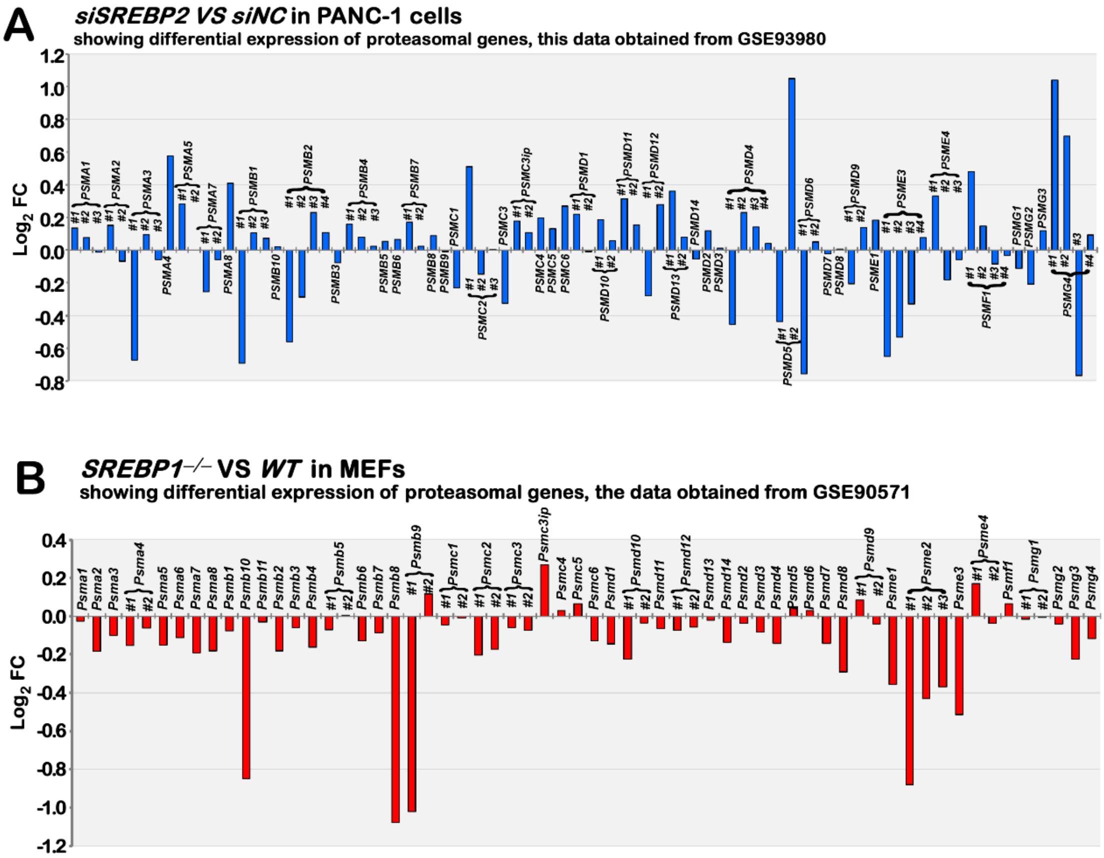


**Figure S3.** Closely scrutinizing two distinct datasets reveal that almost no changes in basal transcriptional expression of all other proteasomal genes except a few of subunits were determined by transcriptomic sequencing of *siSREBP2* (A, from the GSE93980 dataset, https://www.ncbi.nlm.nih.gov/geo/query/acc. cgi? acc=GSE93980) in PANC-1 cells or *SREBP1*$^{-/-}$ MEFs (B, from GSE90571 dataset, https://www.ncbi.nlm.nih.gov/geo/query/acc.cgi?acc= GSE90571).

#### Figure S4

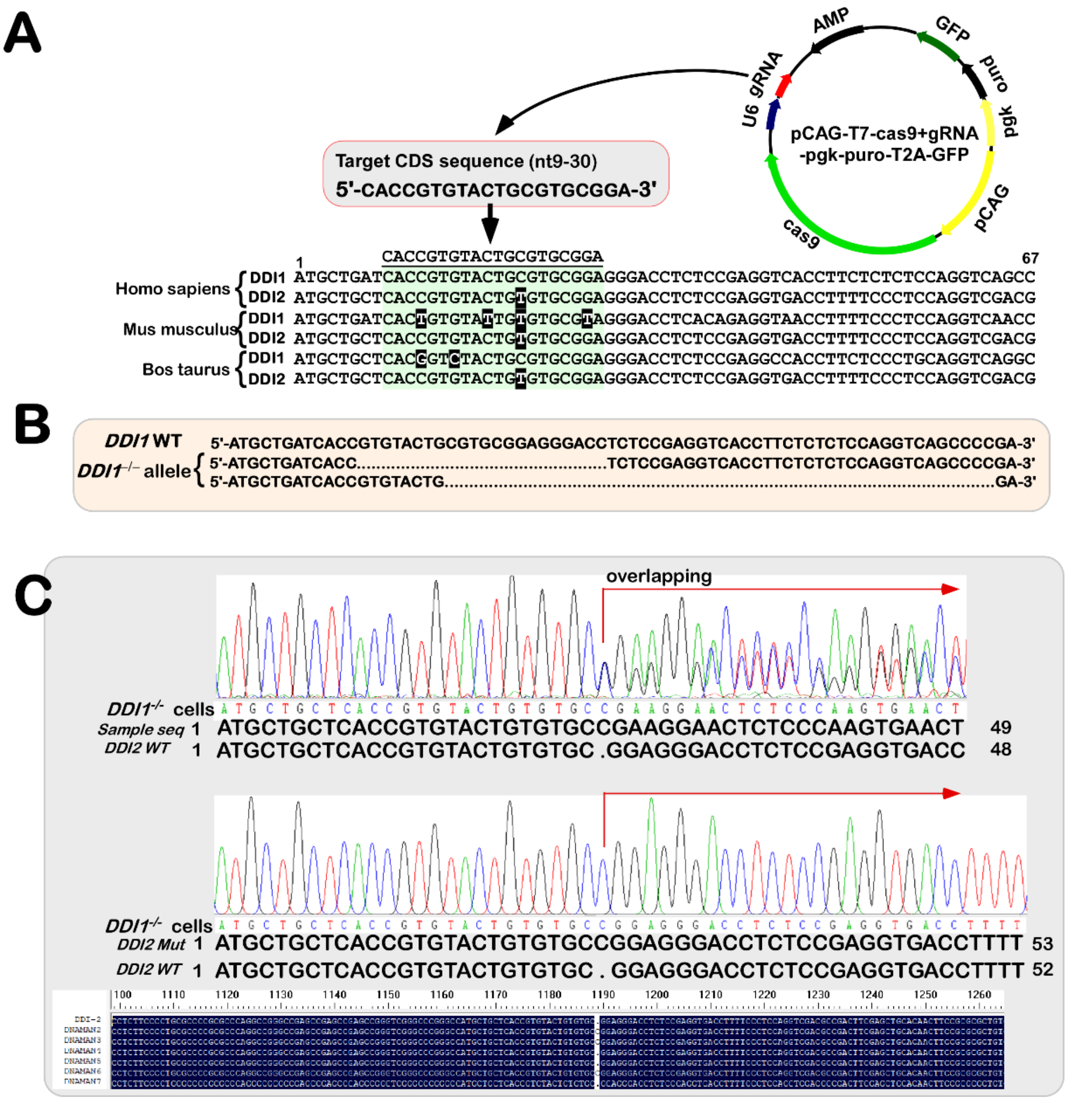


**Figure S4** The human *DDI1*$^{-/-}$ specific gene-editing constructs made by CRISPR/CAS9. (A) Three pairs of nucleotide sequences in CRISPR/CAS9-targeted CDS regions of *DDI1* and *DDI2* from different species are aligned. (B) The resulting knockout mutants (*DDI1*$^{-/-}$) were compared with its wild-type sequence. (C) An extra base of cytosine was inserted in the open-reading frame of *DDI2*, that was identified by the genomic site-specific sequencing in *DDI1*$^{-/-}$ cells; this was collectively designated *DDI1*$^{-/-}$(*DDI2*$^{insC}$), with its location as shown by alignment of the mutant *DDI2*$^{insC}$ with its wildtype nucleotide sequence.

**Table S1. The key reagents and resources used in this study**

| Reagents or resources | Identifier | Source |
|---|---|---|
| **1. Oligonucleotides for qPCR** | | |
| β -actin FW | CATGTACGTTGCTATCCAGGC | Tsingke |
| β -actin REV | CTCCTTAATGTCACGCACGAT | Tsingke |
| Nrf1 FW | TGGAACAGCAGTGGCAAGATCTCA | Tsingke |
| Nrf1 REV | GGCACTGTACAGGATTTCACTTGC | Tsingke |
| Nrf2 FW | TCAGCGACGGAAAGAGTATGA | Tsingke |
| Nrf2 REV | CCACTGGTTTCTGACTGGATGT | Tsingke |
| SREBP1 FW | ACAGTGACTTCCCTGGCCTAT | Tsingke |
| SREBP1 REV | GCATGGACGGGTACATCTTCAA | Tsingke |
| P97 FW | TGGAAACAGATCCTAGCCCTT | Tsingke |
| P97 REV | GCCACCAATGTCATCATACCCT | Tsingke |
| Hrd1 FW | ACCATCTTCATCAAGTATGTGCT | Tsingke |
| Hrd1 REV | TGTACACAGCCTTGTTGTCCC | Tsingke |
| S6K1 FW | ATCGCCACCTGTTCTTACACC | Tsingke |
| S6K1 REV | TCTCCCTCACCTTGCCGACCA | Tsingke |
| DDI-1 FW | ACCACTGTTCCCTGGGCTCCTAC | Tsingke |
| DDI-1 REV | ATGCTGATCACCGTGTACTGCGTGC | Tsingke |
| DDI-2 FW | TCCAGTGCAGTTCCCAAACTTAC | Tsingke |
| DDI-2 REV | ATGCTGCTCACCGTGTACTGTGTGC | Tsingke |
| PSMB5 FW | TCGGCAATGTCGAATCTATGAGC | Tsingke |
| PSMB5 REV | ATGGCTGGGGGCGCAGCGGATTGCA | Tsingke |
| PSMB6 FW | TCAAGAAGGAGGGCAGGTGT | Tsingke |
| PSMB6 REV | AGACTTCTACAACGATCCCCTCT | Tsingke |
| PSMB7 FW | GATACAAGAGCAACTGAAGGGATG | Tsingke |
| PSMB7 REV | ATGGCGGCTGTGTCGGTGTATGCTC | Tsingke |
| **2. Antibodies** | | |
| Nrf1 | Made by us | Zhang's Lab |
| Nrf2 | ab62352 | Abcam |
| p-S6K1 | D151473 | Sangon Biotech |
| β-actin | TA-09 | ZSGB-BIO |
| P97 | ab97302 | Abcam |
| SREBP1 | 14088-1-AP | Proteintech |
| Hrd1 | 13473-1-AP | Proteintech |
| GAPDH | 5174T | CST |
| DDI-1 | H00414301-B01P | Novus Biologicals |
| DDI-2 | A304-630A-T | Bethyl |
| PSMB5 | ab3330 | Abcam |
| PSMB6 | ab150392 | Abcam |
| PSMB7 | ab154745 | Abcam |

| | | |
|---|---|---|
| Keap1 | A1820 | ABclonal |
| **3. Chemicals** | | |
| Rapamycin (RAPA) | 37094 | Sigma Aldrich |
| *tert*-Butylhydroquinone | 112941 | Sigma Aldrich |
| MG132 | M7449 | Sigma Aldrich |
| NAC | A601127 | Sangon Biotech |
| DMSO | 67-68-5 | Aladdin |
| **4. Cell Lines** | | |
| HepG2 | TCHu72 | Cell bank of the Chinese Academy of Sciences |
| HL7702 | GNHu 6 | Cell bank of the Chinese Academy of Sciences |
| *DDI1*$^{-/-}$(*DDI2*$^{insC}$) | Made by us | In this study |
| **5. Oligonucleotides for siRNA** | | |
| siSREBP1 FW | CGGAGAAGCUGCCUAUCAATT | Sangon Biotech |
| siSREBP1 REV | UUGAUAGGCAGCUUCUCCGTT | Sangon Biotech |
| Normal control FW | UUCUCCGAACGUGUCACGUdTdT | Sangon Biotech |
| Normal control REV | ACGUGACACGUUCGGAGAAdTdT | Sangon Biotech |
| **6. Oligonucleotides for expression constructs** | | |
| Nrf1-LUC-#1 FW | CCTAGGCCTGCTAGCGCGACTGAGTTTGTCTCTACACCT | Tsingke |
| Nrf1-LUC-#1 REV | CTTCAGAGAAAAGCTTGCTGAAGGACCAGAATGTTTATGCT | Tsingke |
| Nrf2-LUC FW | CCAGGAGTTTGGTACCAGCCTGGGCAACATAGTGA | Tsingke |
| Nrf2-LUC REV | CCAGCTCCAAGTAGATCTTGATGAGCTGTGGA | Tsingke |
| SREBP1 FW | AGGAGGCGGCCGCGCCATGGACGAGCCACCCTTCA | Sangon Biotech |
| SREBP1 REV | GCTGAGGCCGGGGACTCTAGATCTAGCTGGAAGTGA | Sangon Biotech |
| **7. Recombinant DNAs** | | |
| pcDNA3.1 | V79020 | Invitrogen |
| pGL3-Basic | VQP0121 | Promega |

| | | |
|---|---|---|
| pRL-TK | VQP0126 | Promega |
| **8. Software and Algorithms** | | |
| Canvas X | https://www.canvasgfx.com/ | Canvas GFX, Inc. |
| Chromas 2.4.1 | http://technelysium.com.au/wp/chromas/ | Technelysium Pty Ltd. |
| Excel | https://www.microsoft.com/ | Microsoft |
| Primer Premier 5 | https://www.PremierBiosoft.com/ | PREMIER Biosoft International |
| CFX Manager 3.1 | https://bio-rad-cfx-manager.com/ | Bio-Rad |
| **9. Online data** | | |
| gRNA designed of DDI1 | http://crispr.dbcls.jp/ | CRISPR direct |
| Sequencing expression data in PANC-1 cells | https://www.ncbi.nlm.nih.gov/geo/query/acc.cgi?acc=GSE93980 | GSE93980 dataset |
| Sequencing expression data in MEFs cells | https://www.ncbi.nlm.nih.gov/geo/query/acc.cgi?acc=GSE90571 | GSE90571 dataset |
| **10. Others** | | |
| Cas9/gRNA Construct Kit | VK001 | v-solid |
| Dual-luciferase reporter assay system | E1910 | Promega |
| RNAsimple Total RNA Kit | DP419 | Tiangen Biotech |
| Lipofectamine®3000 Transfection Kit | L3000-015 | Invitrogen |
| GoTaq®qPCR Master Mix | A6001 | Promega |
